# Expanding the Global Map of Protein Post Translational Modifications with Immunoaffinity Enrichment and nDIA Analysis on the Orbitrap Astral Mass Spectrometer

**DOI:** 10.1101/2025.07.03.662636

**Authors:** Mukesh Kumar, Anthony P. Possemato, Barry M. Zee, Srikanth Subramanian, Jian Min Ren, Alissa J. Nelson, Bin Zhang, Sean Landry, Jeffrey C. Silva, Brett Larsen, Tonya Pekar Hart, Matthew P. Stokes, Sean A. Beausoleil

**Author notes:** Correspondence: Mukesh Kumar,; Sean A. Beausoleil.

## Abstract

Post-translational modifications (PTMs) contribute greatly to the diversity of the human proteome by affecting protein structure, function, interactions, stability, localization, and more. The study of PTMs is essential to understand various cellular functions, disease mechanisms, and aid in the development of biomarkers and design of therapeutic targets. Owing to their diversity, dynamic nature, and low stoichiometry compared to unmodified proteome counterparts, the analysis of PTMs remains challenging. In this study, immunoaffinity enrichment of PTM peptides was combined with analysis using Data Dependent Acquisition (DDA) and narrow window Data Independent Acquisition (nDIA) on the Orbitrap Astral mass spectrometer (MS) as well as comparative analysis using the Orbitrap Fusion Lumos MS for Ubiquitination, Phosphorylation, Acetylation, Succinylation, and Methylation. Human cell line and mouse tissue samples at various input peptide amounts were immuno-enriched and mass spectrometry data was acquired on both instruments to assess depth of coverage and number of novel sites identified. The study identified a total of 88,731 unique ubiquitin sites, 64,397 phosphorylation sites (43,721 phosphoserine, 8,414 phosphothreonine and 12,262 phosphotyrosine), 11,629 acetylation, 5,272 succinylation and 1,461 mono-methylation sites. In half the acquisition time, nDIA analysis of immuno-enriched samples on Orbitrap Astral MS provided much greater depth of coverage for all PTMs compared to DDA analysis on Orbitrap Fusion Lumos MS, with up to 33-fold more PTM peptides identified and quantified. Overall, the data presented in this study demonstrates the need for enrichment for PTM detection and the utility of combining antibody-based peptide capture and nDIA on the Orbitrap Astral MS as powerful tools for discovery and profiling of protein post-translational modifications in cells and tissues.

## Introduction

The complexity of the human proteome far surpasses the approximately 20,000 genes that encode it, primarily due to the essential creation of diverse proteoforms. Starting with a limited genomic blueprint, the cellular machinery employs alternative splicing to produce multiple mRNA transcripts from a single gene, each potentially yielding a distinct protein isoform with a unique amino acid sequence. This inherent variability is then massively amplified by a vast array of post-translational modifications (PTMs), such as phosphorylation, acylation, ubiquitination, and methylation, which derivatize these isoforms, altering their structure, localization, interactions, and ultimately their biological roles. The combinatorial effect of alternative splicing and the multitude of possible PTMs results in hundreds of thousands to potentially millions of unique proteoforms within cells and tissues, enabling a dynamic and intricate network of molecular machines that allows for fine-tuned regulation of biological processes and precise responses to a wide range of internal and external cues, far exceeding the information that is directly encoded in the genome [1, 2].

Liquid chromatography (LC) and electrospray ionization tandem mass spectrometry (ESI-MS/MS) are crucial for addressing the challenge of analyzing the proteome, carrying out the identification of endogenously expressed proteins and the detection and localization of a diverse array of post-translational modifications (PTMs) that exist across a wide dynamic range, including those at very low abundance [3-7]. To address the challenge of low abundance PTMs, specialized enrichment strategies targeting specific modifications (e.g., Immobilized Metal Affinity Chromatography (IMAC)-based phosphopeptide enrichment, immunoaffinity-based PTM peptide capture) are often employed prior to LC-MS/MS analysis to increase their detectability [8, 9]. Furthermore, advancements in sample preparation methods that are amenable to automation, robust LC interfaces, mass spectrometry sensitivity as well as data analysis algorithms are continuously improving the ability to identify and quantify more of the proteome, including transient and sub-stoichiometric PTMs within the complex proteomic landscape [10-15].

Immunoaffinity enrichment methods such as PTMScan have aided in overcoming the challenges of monitoring low abundance post-translationally modified peptides and have been used in hundreds of previous studies for the enrichment of PTM peptides prior to LC-MS/MS analysis [9, 16-19][20]. Instead of relying solely on mass spectrometry to detect a minor fraction of PTM peptides or potentially rare, modified peptides within a complex digest, PTMScan utilizes highly specific antibodies that selectively bind to peptides containing a particular PTM or a conserved sequence motif surrounding a modification site [21-24]. These unique antibodies are conjugated to beads, allowing for efficient pull-down and isolation of the modified peptides of interest, effectively increasing their concentration and detectability for downstream LC-MS/MS analysis. This targeted enrichment dramatically reduces the complexity of the sample presented to the mass spectrometer, enabling the identification and quantification of PTMs that might be present at very low stoichiometry and would otherwise be masked by the overwhelming presence of more abundant unmodified peptides in global proteomic approaches.

Traditional data dependent acquisition (DDA) methods of bottom-up proteomics face inherent challenges in comprehensively identifying and quantifying low abundance proteins and low-level post-translational modifications. DDA typically selects the most abundant peptide ions in each mass spectrum for fragmentation and analysis, leading to a sampling bias where less abundant species are often missed. This stochastic selection limits the reproducibility and depth of proteome coverage, particularly for PTMs that are sub-stoichiometric and thus underrepresented. In contrast, next-generation proteomics leverages data independent acquisition (DIA) methods on more advanced mass spectrometers with greater sensitivity and acquisition speed as well as high mass accuracy and resolution to achieve a more complete picture of the sample being analyzed [25-29]. DIA systematically fragments all ions within defined mass-to-charge windows, generating a comprehensive digital map of the biological specimen. This unbiased approach, coupled with the enhanced sensitivity and rapid scanning capabilities of modern instruments, allows for more consistent and deeper proteome coverage, significantly improving the detection and quantification of low abundance proteins and challenging PTMs, even those present at very low levels across a wide dynamic range. The complexity of the resulting DIA data requires sophisticated bioinformatics tools for deconvolution and analysis, but it offers a more thorough perspective of the proteome compared to DDA.

The Thermo Scientific™ Orbitrap™ Astral™ mass spectrometer represents a significant leap forward in mass spectrometry, enabling superior depth and coverage of the proteome in narrow window DIA (nDIA) experiments compared to earlier generation Orbitrap instruments through several key innovations [30, 31]. Firstly, it incorporates a novel mass analyzer using a compact, high-speed design with a 30-meter asymmetric ion track, allowing significantly faster acquisition rates (up to 200 Hz) of high-resolution accurate mass (HRAM) MS/MS spectra with enhanced sensitivity. Secondly, the synchronized parallel acquisition is crucial for nDIA, allowing high dynamic range, high resolution full scan (MS1) acquisition in the Orbitrap analyzer and sensitive HRAM MS/MS data across the entire mass range within the nDIA windows in the Astral analyzer. Thirdly, the improved sensitivity of the Astral analyzer allows for the detection and quantification of lower abundance peptides such as those containing PTMs. This faster acquisition speed, enhanced sensitivity, and parallel analyzer operation on the Orbitrap Astral MS overcome the limitations of earlier Orbitrap instruments in nDIA, providing a much more detailed and quantitative view of the proteome [29-31].

This study illustrates how the combination of immunoaffinity enrichment with Orbitrap Astral MS high sensitivity, speed, and nDIA capabilities represents a leap forward in the ability to comprehensively profile PTMs in cells and tissues. Antibody based affinity enrichment overcomes the challenge of low abundance PTMs by increasing their relative concentration, while the Orbitrap Astral MS nDIA platform provides the depth, sensitivity, and quantitative rigor needed for their comprehensive identification and quantification. nDIA analysis of immuno-enriched samples compared to unenriched samples identified 1,069-fold more ubiquitinated peptides (60 vs. 64,159), 219-fold more phosphotyrosine peptides (43 vs. 9,428), 88-fold more acetylated peptides (127 vs. 11,211), 125-fold more succinylated peptides (46 vs. 5,781) and 19-fold more mono-methylated peptides (74 vs. 1,411). The number of PTM modified peptides identified on the Orbitrap Astral MS using nDIA significantly outpaced the depth of coverage achieved on older generation instruments. For low abundance PTMs like phosphotyrosine, nDIA-HBS analysis on Orbitrap Astral MS identified up to 33-fold more modified peptides compared to DDA analysis on Thermo Scientific™ Orbitrap Fusion™ Lumos™ Tribrid™ mass spectrometer. For high complexity PTMs like ubiquitination, nDIA-HBS on Orbitrap Astral MS identified up to 28-fold more modified peptides compared to DDA on an Orbitrap Fusion Lumos MS

Integrating this detailed PTM information with global proteome profiling data generated on the same advanced instrument will undoubtedly lead to a more complete understanding of the dynamic proteome and the intricate regulatory mechanisms governing cellular processes in response to various stimuli.

## Experimental procedures

### Cell Culture

HCT116 human colorectal carcinoma cells (ATCC CCL-247) were maintained in McCoy’s 5A medium supplemented with 10% fetal bovine serum, 2 mM L-glutamine, 100 U/mL penicillin, and 100 μg/mL streptomycin. Cells were cultured at 37°C in a humidified atmosphere containing 5% CO_₂_ and were passaged when reaching 80-90% confluence using a 0.25% trypsin-EDTA solution. For proteasome inhibition studies, HCT116 cells were seeded in appropriate culture vessels at a density of 5 × 10 cells/mL and allowed to adhere for 24 hours under standard culture conditions. MG-132 (Z-Leu-Leu-Leu-al; Sigma-Aldrich Cat # 474790) was dissolved in DMSO to prepare a 10 mM stock solution, which was stored in aliquots at -20°C. 24 hours after seeding, the culture medium was replaced with fresh complete medium containing 10 μM MG-132 (final DMSO concentration <0.1%). Control cultures received an equivalent volume of DMSO. Cells were incubated for 18 hours at 37°C, 5% CO₂, under standard culture conditions.

Jurkat T cells (ATCC TIB-152) were maintained in RPMI-1640 medium supplemented with 10% fetal bovine serum, 2 mM L-glutamine, 100 U/mL penicillin, and 100 μg/mL streptomycin at 37°C in a humidified atmosphere with 5% CO₂. Cells were passaged every 2-3 days to maintain a density between 0.5-1.5 × 10 cells/mL. For pervanadate treatment, cells in logarithmic growth phase were harvested by centrifugation at 300 × g for 5 minutes at room temperature and resuspended in fresh complete medium at a density of 1 × 10 cells/mL. A 20 mM sodium pervanadate stock solution was freshly prepared immediately before use by combining 1ml 100 mM sodium orthovanadate with 3.4 mL of deionized H2O and 12 uL hydrogen peroxide (30% solution) and incubated at room temperature for 10 minutes. Next, 100 uL of 1M HEPES was added, and the solution was gently mixed. The solution was added to the cell suspension to achieve a final concentration of 1 mM pervanadate. Cells were incubated with pervanadate for 30 minutes at 37°C, 5% CO₂, under standard culture conditions.

Following treatment, cells were washed three times with ice-cold phosphate-buffered saline (PBS), pelleted by centrifugation at 300 × g for 5 minutes at 4°C to remove residual impurities, and flash frozen in a dry ice/ethanol slurry that was prepared by gradually adding 95% laboratory-grade ethanol to crushed dry ice until a semi-fluid mixture was achieved. After the final centrifugation, all supernatant was removed from the cell pellets, and the tubes containing cell pellets were immediately immersed in the dry ice/ethanol slurry for 10 minutes, ensuring tube caps remained above the slurry level. Frozen cell pellets were stored at -80°C until lysis.

### Western blot Analysis

Media was aspirated from culture plates after incubation. Jurkat cells untreated or treated with pervanadate and HCT116 cells untreated or treated with MG-132 were lysed directly in the culture plates by addition of 1× SDS sample buffer containing dithiothreitol (DTT). Cell lysates were scraped from the plates and collected into 1.5 mL microcentrifuge tubes. Samples were sonicated three times for 5 seconds on ice, with 10 seconds rest between each using a probe sonicator at 30% power (Thermo Fisher) to disrupt cellular debris and reduce viscosity, then heated at 95°C for 5 minutes to ensure complete protein denaturation and reduction. Equal volumes of lysates were loaded onto 4-20% gradient polyacrylamide gels (Bio-Rad) and separated by SDS-PAGE at 200V for 40 minutes. Proteins were transferred to nitrocellulose membranes (Bio-Rad) using the Bio-Rad Trans-Blot Turbo System with the mixed molecular weight transfer protocol (2.5A, 25V) for 7 minutes. Transfer efficiency was verified by Ponceau S staining. Membranes were blocked in 5% non-fat dry milk in Tris-buffered saline with 0.1% Tween-20 (TBS-T), shaking for 1 hour at room temperature. Primary antibodies, Phospho-Tyrosine (P-Tyr-1000) MultiMab^®^ Rabbit mAb mix (CST #8954), Ubiquitin (E4I2J) Rabbit mAb (CST #43124), were diluted 1:1000 in 5% bovine serum albumin in TBS-T and incubated overnight with gentle rocking at 4°C. After three washes with TBS-T (5 minutes each), membranes were incubated with horseradish peroxidase-conjugated secondary antibody, Anti-rabbit IgG, HRP-linked Antibody (CST #7074) at a 1:2000 dilution in 5% milk/TBS-T, shaking for 1 hour at room temperature. Following three additional TBS-T washes, proteins were detected using enhanced chemiluminescence reagent SignalFire^™^ ECL Reagent (CST #6883) and visualized with a ChemiDoc imaging system (Bio-Rad).

### Cell / Tissue Lysis, Protein Extraction and Digestion

Human cultured cells and mouse tissue samples were resuspended in 8M Urea, 20mM HEPES pH 8.5, with 2X Phosphatase Inhibitor Cocktail (CST #5870). Resuspended samples were homogenized with a polytron probe homogenizer and sonicated 2 times for 30 seconds at 8 W output power using a microtip with samples placed on ice between each pulse, followed by reduction using DTT (5 mM, 30 minutes) and alkylation using IAA (10 mM, 30 minutes). Samples were diluted to 2 M urea with 20 mM HEPES pH 8.5 with 1 mM CaCl_2_ and digested overnight at 37°C with LysC (Wako-Chem). Samples were further diluted to 1 M urea with 20 mM HEPES pH 8.5 with 1 mM CaCl_2_ and digested for 6 hours with Trypsin (Pierce, Cat# 90058). Samples were acidified with 1:10 volume of 20% trifluoroacetic acid (TFA), and peptides were purified over Waters^TM^ Sep-Pak C18 columns (Cat # WAT023590). An aliquot of the purified peptides was used to determine concentration using a Quantitative Colorimetric Peptide Assay kit (Pierce Cat #23275) and the remainder of the peptide was lyophilized and stored at -80°C.

### Immunoaffinity Enrichment

Immunoaffinity enrichment was carried out using Cell Signaling Technology PTMScan HS kits. The following PTMScan HS kits were used: Ubiquitin/SUMO Remnant Motif (K-ε-GG) Kit (Catalog # 59322); Phospho-Tyrosine (P-Tyr-1000) Kit (Catalog # 38572); Acetyl-Lysine Motif (Ac-K) Kit (Catalog # 46784); Succinyl-Lysine Motif (Succ-K) Kit (Catalog # 60724); Mono-Methyl Arginine Motif (mme-RG) Kit (Catalog # 98567), for the enrichment of ubiquitinated, phosphotyrosine, acetylated, succinylated and mono-methylated peptides. The enrichment was performed according to the protocol provided by the manufacturer. Different input amounts of peptides (0.1, 0.3, 1 and 3 mg) were resuspended in 1.5 mL of 1X HS Immuno Affinity Purification (IAP) buffer. Resuspended peptides were sonicated briefly in a sonicator bath to ensure complete solubilization. The appropriate amount of antibody bead slurry was then added to the peptides followed by end-over-end rotation for 2 hours at 4°C. After binding, the supernatant was transferred to a new Eppendorf tube to allow for sequential enrichment. The antibody bead resin was washed 4 times with 1ml HS IAP wash buffer and 2 times with 1mL water, and peptides were eluted with 50 uL (2x) of 0.15% TFA. Eluted peptides were desalted using C18 tips. The same protocol was repeated for all IAP experiments.

### Immobilized metal affinity chromatography (IMAC) Enrichment

Different peptide inputs were used for IMAC enrichment (0.1 mg, 0.25 mg and 0.5 mg). Peptides were resuspended in 80% MeCN in 0.1%TFA and transferred to a 96 well plate for IMAC enrichment on the AssayMap Bravo (Agilent, CA, USA). Using the provided Phosphopeptide Enrichment application on the Bravo, phospho-peptides were enriched and collected using Fe-NTA cartridges (Agilent, Catalog #G5496-60085). IMAC enriched peptides were resuspended in 0.1% TFA and desalted following the provided Peptide Cleanup 2.0 program on AssayMap Bravo, using 5μl C18 cartridges (Agilent #5190-6532). Clean peptides were dried and resuspended for LC-MS/MS analysis.

### LC-MS/MS Analysis

Prior to LC-MS/MS analysis, peptides were reconstituted in 0.1% FA in 5% MeCN. Peptide mixtures were analyzed by nano LC-MS/MS coupled to either the Orbitrap Astral or Orbitrap Fusion Lumos MS systems. On the Orbitrap Astral MS a Thermo Scientific™ Vanquish™ Neo UHPLC system was used to load samples directly onto an IonOptics C18 Aurora Ultimate column (25 cm x 75 µm ID, 1.7 µm (AUR3-25075C18-TS)). For DIA analysis a 45-minute gradient using Mobile phase A (5% MeCN in 0.1% FA) starting at 2% A and mixing to 30% Mobile phase B (93% MeCN in 0.1% FA) was used at a flowrate of 350 nL/minute. For DDA analysis on the Orbitrap Astral MS a 90-minute gradient using the same buffer start and end points along with the same flowrate was used. For DDA analysis on the Orbitrap Fusion Lumos MS a Thermo Scientific™ EASY-nLC™ 1200 system was used combined with a 50cm analytical column packed in-house with 1.8 µm C18 (Sepax, GP-C18, Catalog # 101181-0000). The same solvents were used as described above to run a 90-minute gradient with the same start and endpoints.

### Mass Spec Parameters

#### Orbitrap Astral MS DDA parameters

MS^1^ spectra were collected in the Orbitrap every 1.2 s at a resolving power of 120,000 at m/z 200 over *m/z* 375–1500 with a standard AGC target and a maximum injection time of 25 ms. The MIPS filter was applied with Peptide mode and “Relax Restrictions when too few Precursors are Found” set to True. Precursors were filtered to charges states 2-6. A Dynamic Exclusion filter was applied with 25 s duration and 10ppm low and high mass tolerance and exclude isotopes set to True. An intensity filter was applied with a minimum precursor intensity of 5000 required for selection. MS^2^ scans were collected in the Astral mass analyzer with an isolation window of 1.6 *m/z*, normalized collision energy of 27, a scan range of 110–2000 *m/z*, a standard AGC target of 100% (1e4 charges), and a maximum injection time of 15 ms.

#### Orbitrap Astral MS nDIA parameters

MS^1^ spectra were collected in the Orbitrap every 0.6 s at a resolving power of 240,000 at m/z 200 over *m/z* 380–980. The MS^1^ normalized AGC target was set to 200% with a maximum injection time of 5 ms. DIA MS^2^ scans were acquired in the Astral analyzer with the Precursor Mass Range set to 380–980 m/z range with the DIA window set to Auto and the Isolation Window set to 2 *m/z.* The Window Placement Optimization was set to off, and the DIA window Mode was set to *m/z* Range. The HCD Collision Energy was set to 25% with the Scan Range in the Astral analyzer set to 150-2000 *m/z* with a Standard AGC Target and an Ion Injection Time set to 3.5 ms.

#### Orbitrap Fusion Lumos MS DDA parameters

MS1 spectra were collected at a resolving power of 120,000 at m/z 200 over *m/z* 300–1500 with a standard AGC target and a maximum injection time of 50 ms. The MIPS filter was applied with Peptide mode selected. Precursors were filtered to charge states of 2-7 and a Dynamic Exclusion filter was applied with 30 s duration and 10ppm low and high mass tolerance. MS^2^ scans were collected in the Orbitrap analyzer at a resolution of 30,000 at m/z 200 with a defined first mass of 110 *m/z* and an isolation window of 1.6 *m/z.* A fixed Collision Energy of 27 was used with a Standard AGC and a maximum injection time of 50 ms.

### Data Processing

DIA mass spectra were processed using Spectronaut® v19 (Biognosys AG, Switzerland) [32]. DirectDIA® module was used for searching using the default parameters with the following changes: Fixed modification was set to Carbamidomethyl (C) and specific variable modifications were added for different enrichments: GlyGly (K) for ubiquitination, Acetyl (K) for Acetylation, Methyl (R) for Mono-Methylation, Phospho (STY) for phosphorylation, and Succinyl (K) for Succinylation. Minimum peptide length 6, Missed Cleavage 3 and Max Variable Modification was set to 5. For Identification, PSM FDR -0.01, Peptide FDR -0.01, Protein Group FDR -0.01, PTM Localization Filter Checked with Min Localization Threshold 0.75. In Result Filters, Best N Fragments per peptide Max 20 and Min 3. Peptide charge (Max 4, and Min 2). For DIA Hybrid search, all parameters remain the same as DIA LFS except for checking the box Hybrid (DDA + DIA) Library in the workflow section. After searching, the standard peptide report as well as a PTM site report was exported and further processed. To determine the number of modified peptides, data were filtered by “EG.ModifiedSequence”, then sorted by “PEP.RunEvidenceCount” (modified peptides were only retained with minimum of 1 for PEP.RunEvidenceCount). Next, among the retained modified peptides, duplicate entries representing multiple precursors were deleted and only unique modified sequences were retained. Next, the retained unique modified sequence was sorted according to “EG.TotalQuantity” and any peptide with a zero value was omitted from the list of identified peptides.

DDA data was processed using MSFragger v4 [33]. Default parameters were used with the following changes: For the ion quant MaxLFQ, Match between runs and normalize intensity across runs were unchecked. As with DIA, search specific modifications were added. For the analysis of modified peptides, the “combined_modified_peptide.tsv” file was used. First the peptide was filtered based on “modified sequence” column containing the specific modifications. Next the modified peptides were sorted according to “Intensity” and peptides with zero intensity not included in the analysis.

### Experimental Design and Statistical Rationale

The study aimed to evaluate the combined use of immunoaffinity enrichment, and the nDIA analysis on the Orbitrap Astral MS for the identification and quantification of PTM peptides. The experiment was designed to enrich PTM peptides from four different amounts of input peptide (0.1 mg, 0.3 mg, 1.0 mg and 3.0 mg) and samples from each input amount were analyzed by DDA and nDIA analysis on Orbitrap Astral MS as well as by DDA on Orbitrap Fusion Lumos MS. Analyses of different PTMs including Phosphorylation, Ubiquitination, Acetylation, Succinylation and Mono-Methylation are presented in the study. The need for enrichment was tested by running unenriched samples in parallel on the Orbitrap Astral MS with nDIA and searching for each PTM.

## Results

### Overview of the workflow used for the enrichment and analysis of PTM peptides

To evaluate the efficacy of immunoaffinity enrichment and the Orbitrap Astral MS for the analysis of PTMs, proteins from untreated and treated human cell lines (HCT116, Jurkat) and mouse liver tissue were digested to peptides and enriched with the appropriate PTM antibody (Ubiquitin, Tyrosine phosphorylation, Acetylation, Methylation, or Succinylation) or Fe-IMAC beads (Serine/Threonine phosphorylation). Enriched peptides were analyzed using Data Dependent Acquisition (DDA) on the Orbitrap Fusion Lumos MS and by DDA and narrow window Data Independent Acquisition (nDIA) on the Orbitrap Astral MS. Acquired mass spectrometric DDA and nDIA data were processed using MSFragger and Spectronaut, respectively. A schematic of the workflow used in this study is shown in **Figure 1**.

**Figure 1:**
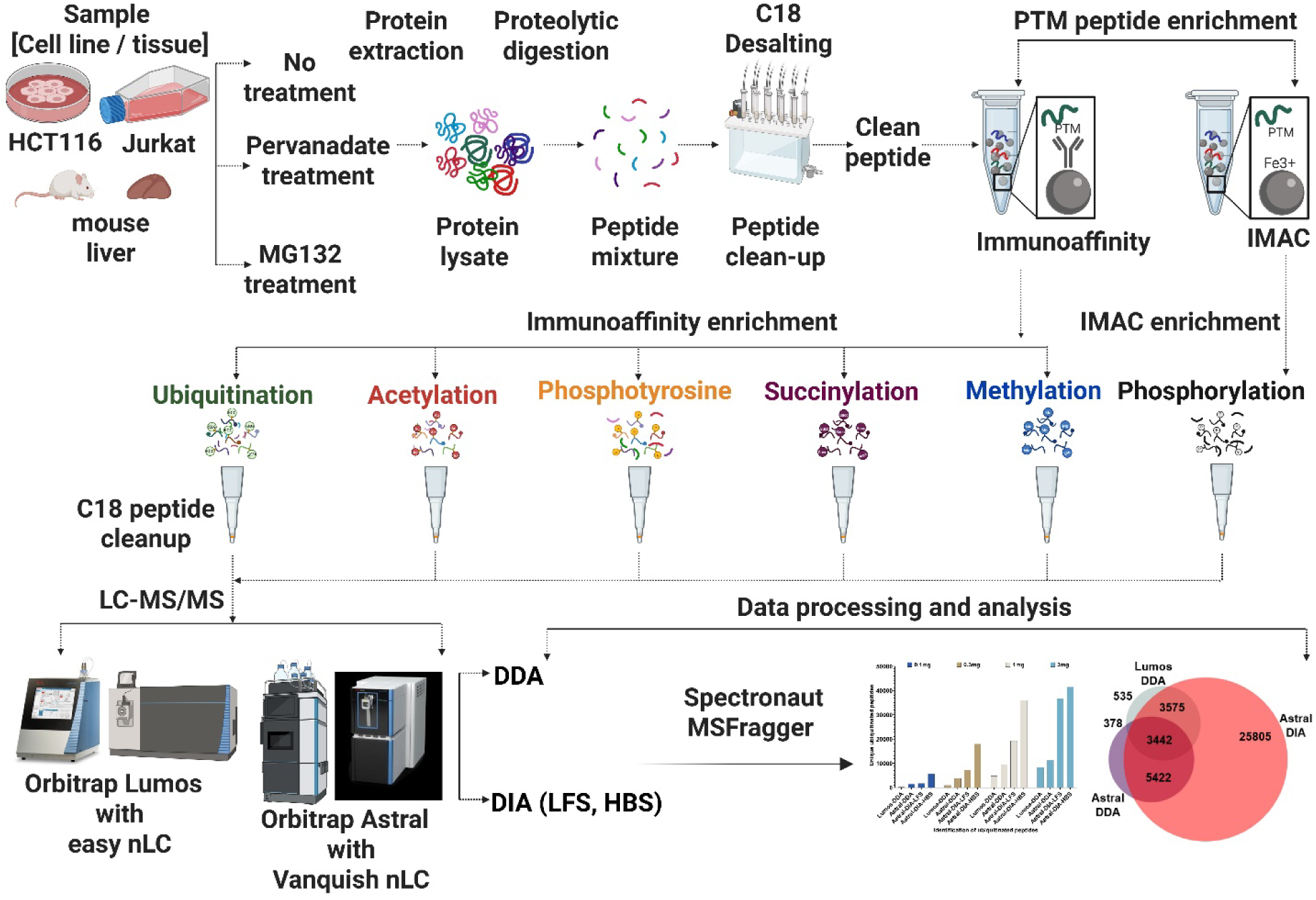
Schematic of the workflow used for the enrichment and analysis of PTM peptides. Total proteins were extracted from untreated or MG132 treated human colorectal carcinoma (HCT116) cells, untreated or pervanadate treated T-cell acute lymphoblastic leukemia (Jurkat) cells, and from untreated mouse liver tissue. Protein extracts were reduced, alkylated and digested to peptides using trypsin and Lys-C. Digested peptides were desalted using Sep-Pak C18 cartridges. Peptides were enriched with Fe-IMAC for serine/threonine/tyrosine phosphopeptides. PTMScan HS kits were used to enrich ubiquitin remnant (K-ε-GG), lysine-acetylation, tyrosine-phosphorylation, lysine-succinylation, and mono-methylation of arginine. Prior to LC-MS/MS analysis, enriched peptides were desalted using C18 tips, loaded onto a reverse phase C18 column, and separated at a nanoflow rate using Easy-nLC system or Vanquish Neo UHPLC system connected to Orbitrap Fusion Lumos and Orbitrap Astral mass spectrometers, respectively. Data was acquired using data dependent acquisition (DDA) and narrow window data independent acquisition (nDIA), and processing of the data was carried out using Spectronaut and MSFragger.

### Immunoaffinity Enrichment and Analysis of Ubiquitinated Peptides

To evaluate the performance Orbitrap Astral MS for the analysis of ubiquitination, cellular proteins from the human colorectal carcinoma HCT116 cell line were extracted and digested to peptides. Different amounts of total peptide input (0.1 mg, 0.3 mg, 1 mg and 3 mg) were used for immuno-enrichment of ubiquitin remnant-containing peptides using the PTMScan® HS Ubiquitin/SUMO Remnant Motif (K-ε-GG) Kit (Catalog # 59322). A ubiquitin K-ε-GG containing peptide retains the diglycine remnant from ubiquitin after proteolytic digestion by trypsin [34]. This remnant, a modified lysine residue (K-ε-GG), serves as a signature of ubiquitination, allowing identification of specific ubiquitinated sites on proteins. Enriched peptides were analyzed by DDA on Orbitrap Fusion Lumos MS and by DDA and nDIA on Orbitrap Astral MS. From the 3 mg input amount, DDA analysis on Orbitrap Fusion Lumos MS and Orbitrap Astral MS identified 8,391 and 11,244 unique ubiquitinated peptides (K-ε-GG), compared to 36,691 and 41,469 unique ubiquitinated peptides (K-ε-GG) by nDIA-LFS and nDIA-HBS analysis respectively (**Figure 2A**). These unique peptides mapped to 2,789, 3,564, 6,288 and 6,441 unique protein groups, respectively. In half the mass spec acquisition time (45-minute vs. 90-minute), nDIA analysis on Orbitrap Astral MS identified up to 4.9- and 3.6-fold more unique ubiquitinated peptides compared to DDA analysis on Orbitrap Fusion Lumos MS and DDA analysis on Orbitrap Astral MS, respectively. From the lowest input amount (0.1 mg) 204, 1379, 1716, and 5724 unique ubiquitinated peptides were identified using Lumos-DDA, Astral-DDA, Astral-DIA-LFS and Astral-DIA-HBS, respectively. The Astral-DIA-HBS analysis identified up to 28.0-, 4.0- and 3.0-fold more ubiquitinated peptide compared to Lumos-DDA, Astral-DDA and Astral-DIA-LFS analysis, respectively (**Figure 2A**). When using the hybrid search (HBS) option in Spectronaut, nDIA and DDA data from all input amounts were added in the search as library extension runs. When comparing with nDIA-LFS, the benefit of HBS was more pronounced at the lower input amount and the benefit was minimized at the highest input amount as shown by the 3.3-, 2.4-, 1.8- and 1.1-fold increase in the identification of ubiquitinated peptides from 0.1 mg, 0.3 mg, 1 mg and 3 mg, respectively. nDIA analysis on Orbitrap Astral MS yielded >25,000 additional unique peptides when compared with DDA analysis on Orbitrap Fusion Lumos MS and Orbitrap Astral MS (**Figure 2B**). To further validate and test the extent of the number of ubiquitinated peptides that can be identified by Orbitrap Astral MS, HCT116 cells were treated with the proteasome inhibitor MG132 to maximize the presence of ubiquitinated proteins in the sample. Western blotting of whole cell lysates from untreated and MG132 treated cells using anti-ubiquitin antibody confirmed the increased level of ubiquitinated proteins in MG132 treated cells compared to untreated cells (**Figure 2C**).

**Figure 2:**
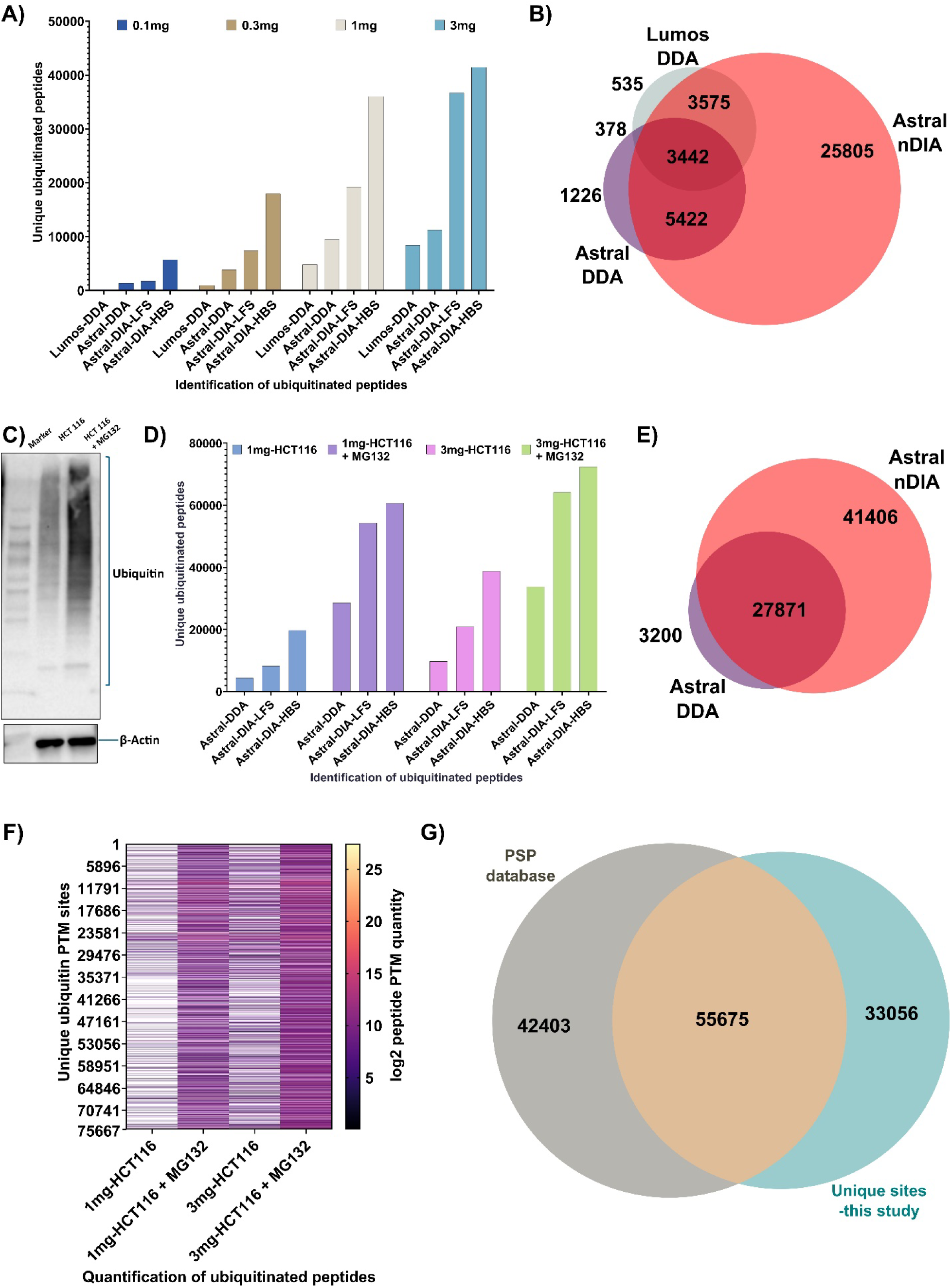
Enrichment and analysis of ubiquitinated peptides. **A)** The number of unique ubiquitinated peptides identified and quantified in cells by Lumos DDA, Astral DDA, Astral nDIA-LFS (Library Free Search), and Astral nDIA-HBS (Hybrid search), from different amounts of input peptide used for immunoaffinity enrichment. **B**) The overlap of ubiquitinated peptides identified and quantified from 3 mg input across different acquisition methods. **C**) Anti-ubiquitin western blot of untreated and MG132 treated HCT116 cells. **D**) The number of unique ubiquitinated peptides identified and quantified by Astral-DDA, Astral-nDIA-LFS, and Astral-nDIA-HBS from untreated and MG132 treated HCT116 cells at different input peptide amounts used for immunoaffinity enrichment. **E**) The overlap of ubiquitinated peptides identified and quantified by Orbitrap Astral MS DDA and nDIA acquisition methods. **F**) The quantification of ubiquitinated PTM peptides identified by Orbitrap Astral DIA from untreated and MG132 treated HCT116 cells. **G**) The overlap of ubiquitin sites identified in this study when compared to all ubiquitin sites reported in the PhosphoSitePlus (PSP) database [35].

1 mg and 3 mg input peptide from untreated and MG132 treated HCT116 cell line samples were used for immunoaffinity enrichment for ubiquitinated peptides and analyzed by DDA and nDIA analysis on Orbitrap Astral MS. From the 3 mg input amount more than 72,000 unique ubiquitinated peptides were identified from MG132 treated cells, 6-fold more than in untreated cells (**Figure 2D**). When comparing the number of ubiquitinated peptides identified by DDA and nDIA analysis, a total of 27,871 were identified in both DDA and nDIA runs, with 3,200 peptides identified uniquely in DDA, and 41,406 peptides uniquely identified in nDIA (**Figure 2E**). These numbers were achieved using a shorter, 45-minute gradient for nDIA and a longer, 90-minute gradient for DDA. Quantification of PTM sites confirmed the increased level of ubiquitination in MG132 treated cells compared to untreated cells (**Figure 2F**). Out of 88,731 unique ubiquitin sites identified in this study, a total of 55,675 sites matched to known sites when mapped against the PhosphoSitePlus® database [35] and a total of 33,056 sites are uniquely identified in this study (**Figure 2G**). Identification and quantification of all ubiquitinated peptides by DDA and nDIA are provided in **Supplementary Data S1A and S1B**, respectively.

### Enrichment and Analysis of Phosphorylated Peptides

Immobilized Metal Affinity Chromatography (IMAC) was used as a general enrichment tool for phosphorylated peptides. 0.1 mg, 0.25 mg, and 0.5 mg of HCT116 peptides were used for IMAC enrichment and analyzed by LC-MS/MS using DDA acquisition on Orbitrap Fusion Lumos MS, and DDA and nDIA analysis on Orbitrap Astral MS. From the 0.1 mg input amount, a total of 13,654, 21,870, 19,206 and 28,275 unique phosphorylated peptides were identified by Lumos-DDA, Astral-DDA, Astral-nDIA-LFS and Astral-nDIA-HBS, respectively (**Figure 3A**). A similar number of phosphorylated peptides were identified in both the 0.25 mg and 0.5 mg peptide input amount for the IMAC enrichment 17,304 (0.25 mg), 17,271 (0.5 mg); 24,170 (0.25 mg), 24,187 (0.5 mg); 23,150 (0.25 mg), 22,993 (0.5 mg), 32,339 (0.25 mg), and 32,480 (0.5 mg) were identified by Lumos-DDA, Astral-DDA, Astral-nDIA-LFS and Astral-nDIA-HBS, respectively (**Figure 3A**).

**Figure 3:**
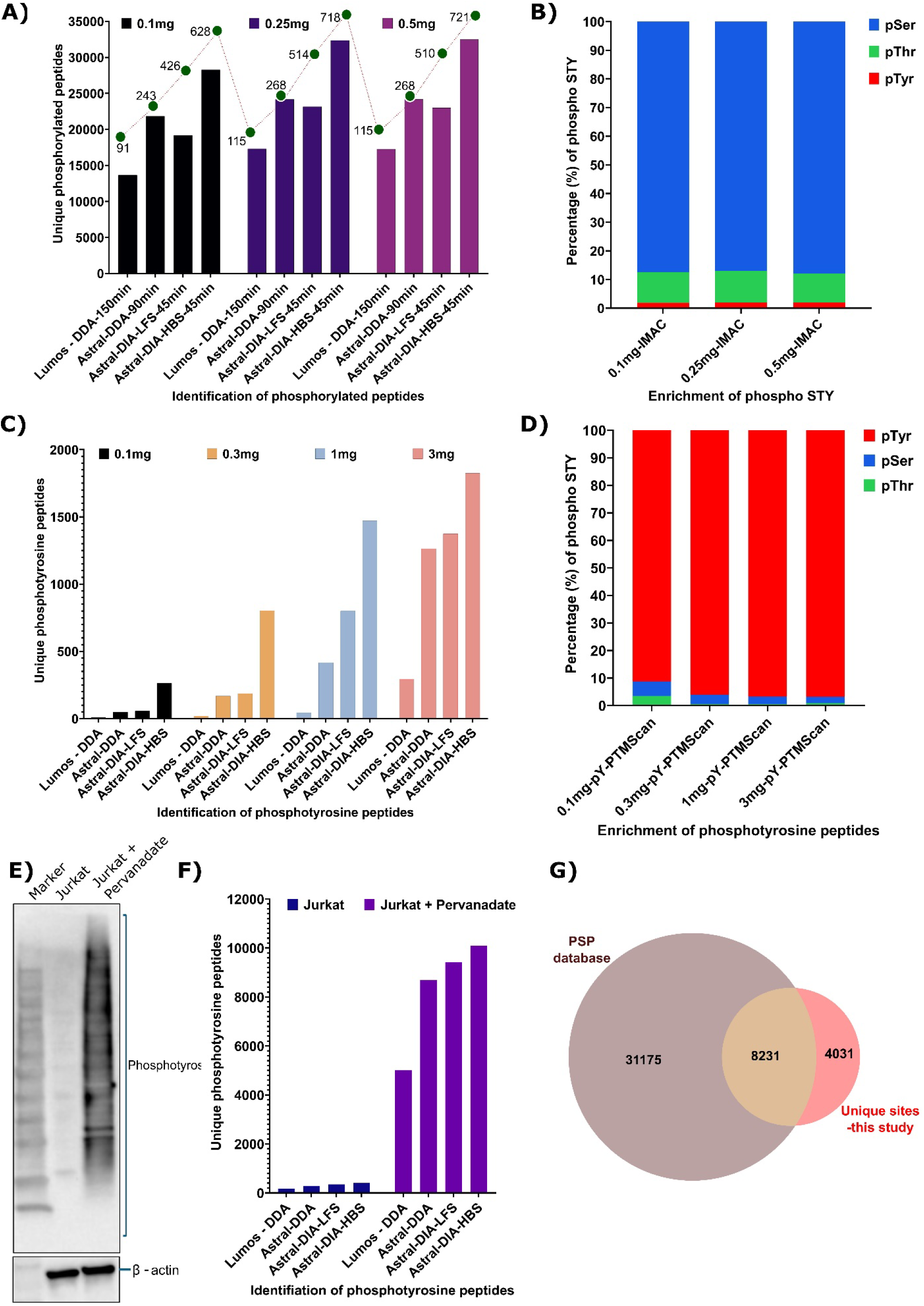
Enrichment and analysis of phosphorylated peptides. **A)** The number of unique phosphorylated peptides identified and quantified by Lumos-DDA, Astral-DDA, Astral-DIA-LFS (Library Free Search), and Astral-DIA-HBS (Hybrid search), from different input peptide amounts used for IMAC enrichment. The number on top of each bar represents the number of quantified phosphopeptides per minute by the respective methods. **B**) The percentage of phosphoserine (pSer), phosphothreonine (pThr) and phosphotyrosine (pTyr) peptides identified and quantified by IMAC enrichment at different input amounts. **C**) The number of unique phosphotyrosine peptides identified and quantified by Lumos-DDA, Astral-DDA, Astral-nDIA-LFS, and Astral-nDIA-HBS from different peptide input amounts used for immunoaffinity phosphotyrosine enrichment. **D**) The percentage of pSer, pThr and pTyr peptides identified and quantified by immunoaffinity phosphotyrosine enrichment at different input amounts. **E**) An anti-phosphotyrosine western blot of untreated and pervanadate (a protein tyrosine phosphatase inhibitor) treated Jurkat cells. **F**) The number of unique phosphotyrosine peptides enriched by immunoaffinity and identified and quantified by Lumos-DDA, Astral-DDA, Astral-nDIA-LFS, and Astral-nDIA-HBS from untreated and pervanadate treated Jurkat cells. **G**) The overlap of phosphotyrosine sites identified in this study when compared to all phosphotyrosine sites reported in the PhosphoSitePlus® database.

nDIA analysis of IMAC enriched samples on the Orbitrap Astral MS identified up to 6-fold more phosphorylated peptides per minute compared to the Orbitrap Fusion Lumos MS at each of the input amounts tested (**Figure 3A**). Among all phosphorylated peptides detected, approx. 87% were serine phosphorylated, 11% threonine phosphorylated, and approx. 2% tyrosine phosphorylated (**Figure 3B**). To overcome the limitation of IMAC for the enrichment of phosphotyrosine, immunoaffinity was used to enrich phosphotyrosine-containing peptides. 0.1 mg, 0.3 mg, 1 mg, and 3 mg of input peptides from HCT116 cells were used to evaluate enrichment efficacy of immunoaffinity capture and the analytical performance of Orbitrap Astral MS for detection of low abundance phosphotyrosine peptides. From the lowest input amount used for enrichment (0.1 mg), Orbitrap Astral MS DDA, nDIA-LFS and nDIA-HBS identified 6x, 7x and 32x more phosphotyrosine peptides respectively compared to Orbitrap Fusion Lumos MS DDA (**Figure 3C**). From the 3 mg input amount nDIA on Orbitrap Astral MS identified 1,824 phosphotyrosine peptides compared to 294 phosphotyrosine peptides by Orbitrap Fusion Lumos MS DDA (**Figure 3C**). In contrast to IMAC, 97% of all phosphorylated peptides enriched by immunoaffinity were tyrosine phosphorylated, with the remaining approx. 2% serine phosphorylation, and approx. 1% threonine phosphorylation (**Figure 3D**). To further validate, Jurkat cells were treated with pervanadate - a potent protein tyrosine phosphatase (PTP) inhibitor commonly used to increase tyrosine phosphorylation in cells. Western blotting of whole cell lysate from untreated or pervanadate treated Jurkat cells using a pan anti-phosphotyrosine antibody confirmed the increased level of protein tyrosine phosphorylation with pervanadate treatment (**Figure 3E**). Peptides from untreated and pervanadate treated Jurkat cells were used for immunoaffinity based phosphotyrosine enrichment followed by LC-MS/MS analysis. 8,687, 9,428, and 10,099 unique peptides containing tyrosine phosphorylation were detected by DDA, nDIA-LFS and nDIA-HBS analysis on Orbitrap Astral MS, compared to 5,007 phosphotyrosine peptides from DDA analysis on the Lumos (**Figure 3F**). Pervanadate treatment of samples increased the number of phosphotyrosine peptides identified 31-fold (167 vs. 5,007), 30-fold (282 vs. 8,687), 27-fold (349 vs. 9,428) and 25-fold (407 vs. 10,099) with Lumos DDA, Astral DDA, Astral-nDIA-LFS and Astral-nDIA-HBS respectively (**Figure 3F**), in line with the analysis by Western blot (**Figure 3E**). Identification and quantification of all phosphorylated peptides by DDA and nDIA are provided in **Supplementary Data S2A and S2B**, respectively. When mapped against the PhosphoSitePlus database, out of 12,262 phosphotyrosine sites identified, 8,231 sites matched to previously identified sites, and 4,031 phosphotyrosine sites were uniquely identified in this study (**Figure 3G**).

### Enrichment and Analysis of Acetylated, Succinylated and Mono-methylated Peptides

To further test the application of immunoaffinity enrichment and nDIA on the Orbitrap Astral MS, the analysis was extended to other biologically important PTMs such as acetylation and succinylation of lysine and mono-methylation of arginine. For acetylation analysis, 0.1 mg, 0.3 mg, 1 mg and 3 mg of total peptide from HCT116 cells were used for immunoaffinity enrichment followed by LC-MS/MS analysis. From the lowest input amount of 0.1 mg peptide, a total of 576, 857, 870 and 2,954 unique acetylated peptides were identified by Lumos DDA, Astral DDA, Astral nDIA-LFS and Astral nDIA-HBS respectively (**Figure 4A**). From the 3 mg input amount, Orbitrap Astral MS nDIA-HBS identified a total of 13,142 unique acetylated peptides, 2.7-fold more than the 4,743 acetylated peptides identified by Orbitrap Fusion Lumos MS DDA (**Figure 4A**). nDIA on Orbitrap Astral MS identified a similar number of unique acetylated peptides as DDA analysis in half the run time (45-minute vs. 90-minute gradient) (**Figure 4A**). Out of the 11,629 unique acetylated sites identified in this study, 5,682 sites were previously identified in the PhosphoSitePlus database, and a total of 5,947 sites were uniquely identified in this study (**Figure 4B**). Tryptic peptides from mouse liver were used for the enrichment of succinylated and arginine mono-methylated peptides. 1 mg and 3 mg input peptide were used for the immunoaffinity enrichment followed by LC-MS/MS analysis.

**Figure 4:**
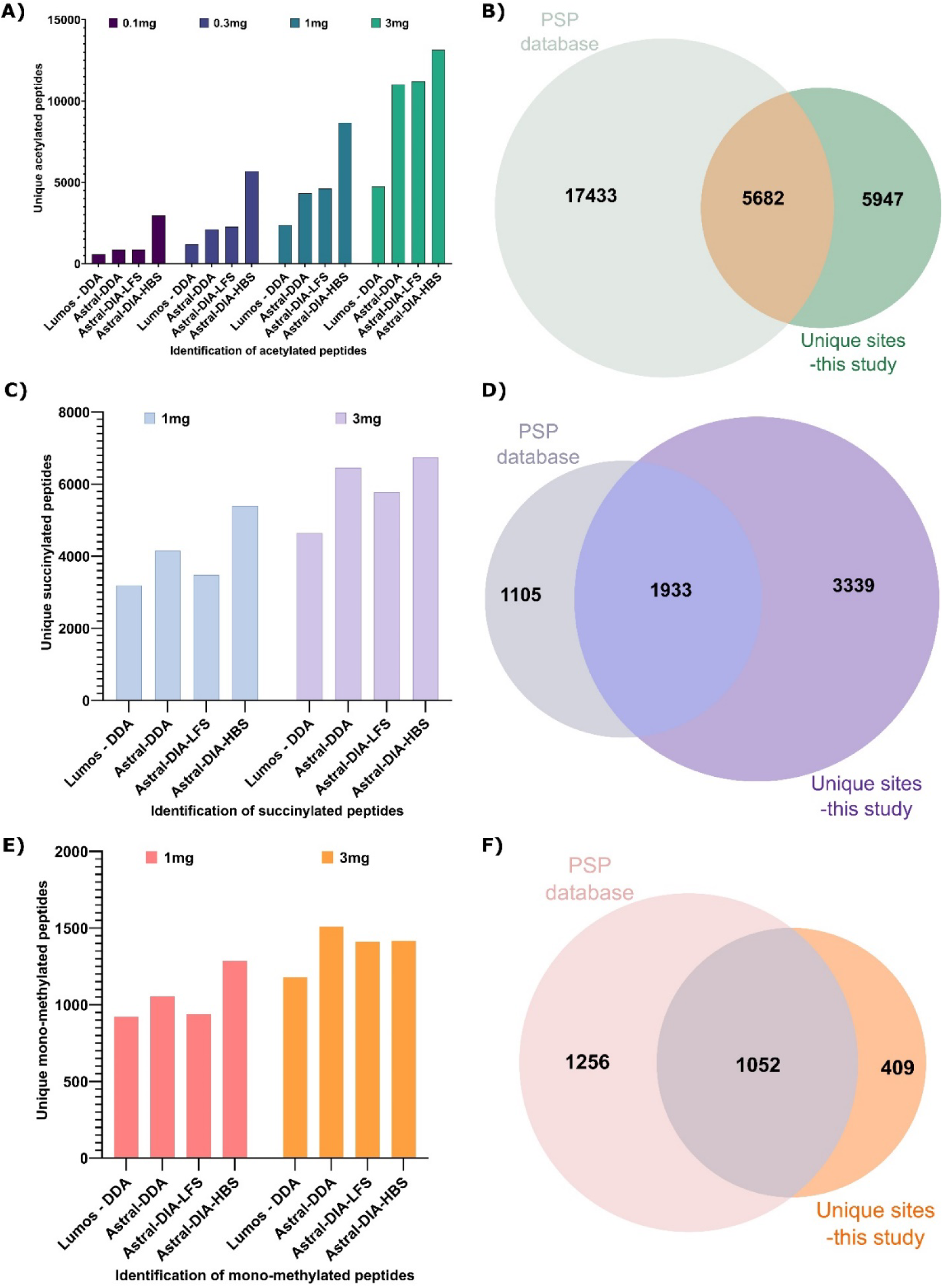
Enrichment and analysis of acetylation, succinylation and arginine methylation. **A, C, E)** Panels A, C and E, respectively show the number of unique acetylated, succinylated and arginine mono-methylated peptides identified and quantified by Lumos DDA, Astral DDA, Astral DIA-LFS, and Astral DIA-HBS from different amounts of input peptide used for immunoaffinity enrichment. **B, D, F**) Panels B, D and F show the overlap of acetylation, succinylation and arginine mono-methylation sites identified in this study when compared to all acetylation, succinylation and arginine mono-methylation sites reported in the PhosphositePlus® (PSP) database, respectively.

nDIA HBS analysis on Orbitrap Astral MS identified 5,401 and 6,742 unique succinylated peptides from 1 mg and 3 mg input amount, compared to 3,193 and 4,640 succinylated peptides, respectively, by Orbitrap Fusion Lumos MS DDA (**Figure 4C**). When compared to nDIA-LFS, DDA analysis on Orbitrap Astral MS using a 2x longer gradient (90-minute vs. 45-minute), identified 20% and 11% more succinylated peptides from 1 mg and 3 mg respectively. When the same nDIA data are searched in hybrid mode (DIA-HBS) using DDA runs to extend the library, nDIA-HBS identified up to 30% more succinylated peptides compared to DDA in half the analysis time. When mapped against the PhosphoSitePlus database, 1,933 succinylated sites were previously known and 3,339 sites are uniquely identified in this study (**Figure 4D**). immunoaffinity analysis of arginine mono-methylation from mouse liver resulted in the identification of 1,179, 1,508, 1,411, and 1,417 unique arginine mono-methylated peptides by Lumos-DDA, Astral-DDA, Astral-nDIA-LFS and Astral-nDIA-HBS, respectively from 3 mg input peptide (**Figure 4E**). From the 1 mg input enrichment, Astral-nDIA-HBS identified 39%, 21% and 36% more arginine monomethylated peptide compared to Lumos-DDA, Astral-DDA and Astral-nDIA-LFS, respectively (**Figure 4E**). Comparing to the PhosphoSitePlus database, 1,052 arginine mono-methylation sites were previously identified in other studies, and 409 sites were uniquely identified in this study (**Figure 4F**). Identification and quantification of all acetylated, succinylated and mono-methylated peptides by DDA and nDIA are provided in **Supplementary Data S3A and S3B**, respectively.

### Need for Enrichment of PTM Peptides Prior to LC-MS/MS analysis

Technological advancements like the Orbitrap Astral MS have provided unprecedented speed and sensitivity and have enabled the detection of tens to hundreds of thousands of peptides in a short analysis time, providing in-depth analysis of thousands of protein groups in a typical bottom-up proteomics analysis. Even with this increased speed and sensitivity, the detection of PTM peptides in unenriched HCT116 cells from a 45-minute nDIA-LFS method quantified a total of 112,440 unique peptides from 8,440 protein groups but only 24 (0.02%) ubiquitinated, 127 (0.1%) acetylated, and 6 phosphotyrosine (0.005%), peptides were identified (**Supplementary Data S4**). However, when immunoaffinity enrichment methods such as PTMScan HS were employed on HCT116 cells prior to nDIA-LFS analysis, a total of 36,691 unique ubiquitinated sites were detected (**Figure 5A**). Even in MG132-treated HCT116 cells, only 60 ubiquitinated peptides were identified in unenriched samples compared to 64,159 ubiquitinated peptides with immunoaffinity enrichment (**Figure 5B**). A total of 1,374 phosphotyrosine peptides were identified with immunoaffinity enrichment vs. 6 phosphotyrosine peptides in unenriched HCT116 samples (**Figure 5C**).

**Figure 5:**
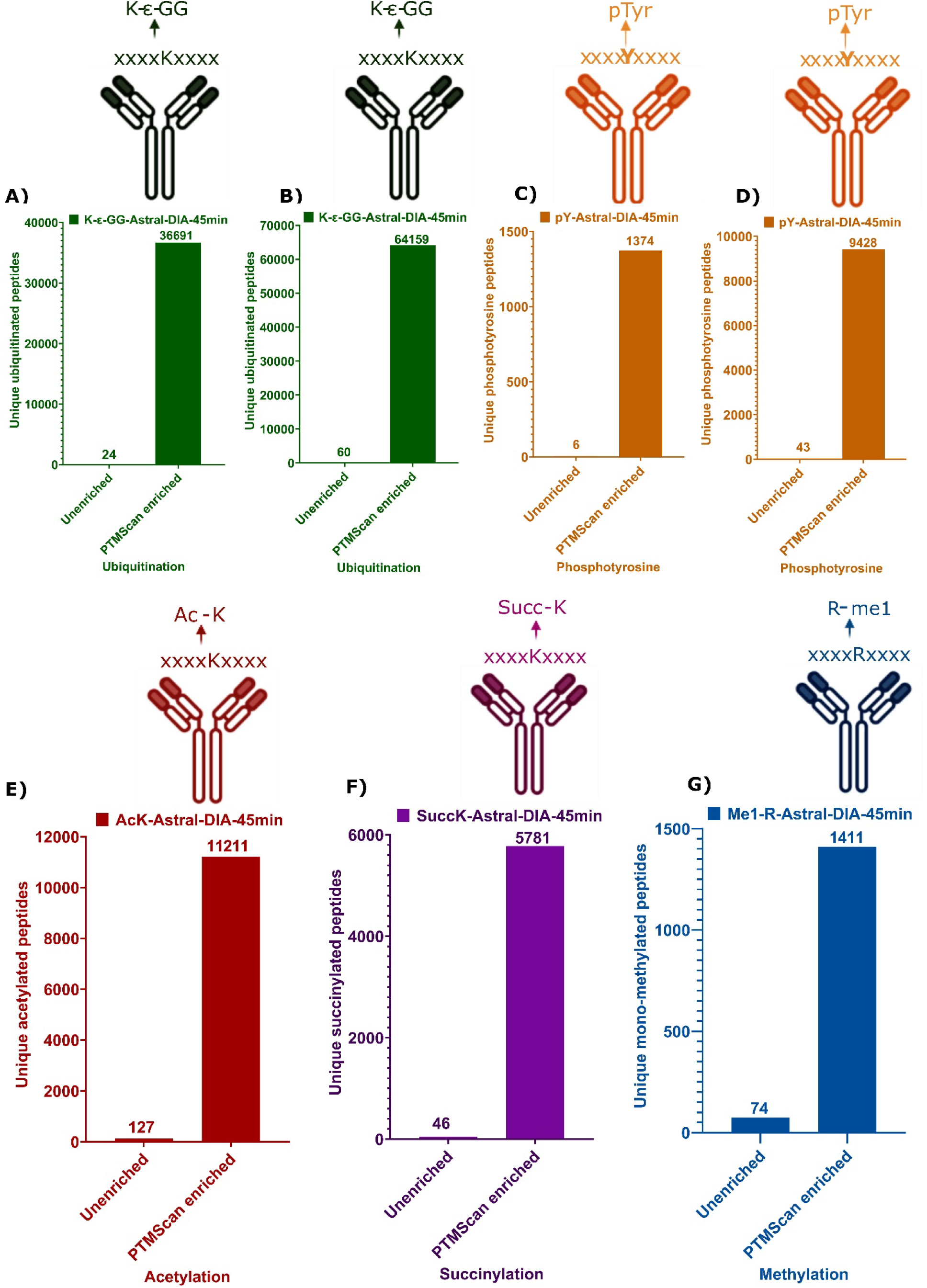
Comparison of PTM site identification with and without enrichment. Panel A – G shows the number of quantified PTM sites by single shot library free DIA analysis on Orbitrap Astral MS from an unenriched and corresponding PTMScan enriched sample.

In Jurkat cells treated with pervanadate, where the phosphotyrosine level is approx. 28-fold higher than untreated cells (**Figure 3E**, **3F**), only 43 phosphotyrosine peptides were identified without enrichment compared to 9,428 phosphotyrosine peptides with immunoaffinity enrichment (**Figure 5C**). Similarly, the number of acetylated, succinylated and monomethylated peptides showed a similar pattern of low identification rate in unenriched samples versus more comprehensive coverage in immunoaffinity enriched samples 127 vs. 11,211 for acetylation, 46 vs. 5,781 for succinylation, and 74 vs. 1,411 for arginine mono-methylation **Figure 5E**, **Figure 5F and Figure 5G**, respectively.

## Discussion

The study of protein post translational modifications is critical to understanding the signaling and protein dynamics underlying cellular processes and disease states. IMAC and antibody based enrichment methods such as PTMScan have long be used to enrich for post translationally modified peptides prior to LC-MS/MS analysis, simplifying the mixture of peptides delivered to the instrument and allowing identification and quantification of thousands of sites of modification across a variety of PTMs [9, 16-19]. Recent advancements in instrumentation and software have allowed unprecedented coverage of the proteome, nearing a comprehensive view of protein expression [28, 30, 31], however we wanted to understand how new instruments like the Orbitrap Astral MS could further improve the coverage of PTMs with or without conventional enrichment methods typically employed in LC-MS based PTM studies. In this study the performance of the Orbitrap Astral MS instrument using both DDA, library-free nDIA, and hybrid search nDIA was compared to an older generation Orbitrap Fusion Lumos tribrid MS instrument on unenriched and a variety of enrichment samples to assess depth of coverage on multiple PTM types and the ability to uncover novel sites (**Figure 1**).

Lysine ubiquitination plays a central role in protein degradation as well as regulating cell signaling events [36-40]. The PTMScan Ubiquitin remnant enrichment kit recognizes the di-glycine motif left behind after trypsin digestion of ubiquitinated sites (K-ε-GG). Using this antibody to enrich samples previously allowed identification of thousands of sites of ubiquitination on older generation instruments such as the Orbitrap Fusion Lumos tribrid MS. In this study, the number of identified and quantified ubiquitin sites increased across all sample amounts (0.1 mg, 0.3 mg, 1 mg, 3 mg), with nDIA enabling much larger gains in number of identified peptides, up to a 28-fold increase compared to Orbitrap Fusion Lumos DDA (**Figure 2A**, **2B**). The proteasome inhibitor MG132 was used to provide an even richer sample from which to enrich ubiquitinated peptides, and at both 1 mg and 3 mg input peptide the number of ubiquitin K-GG peptides identified increased compared to untreated control, up to nearly 80,000 unique peptides in the 3 mg sample on Orbitrap Astral MS with DIA-HBS (**Figure 2C**, **2D**, **2E**, **2F**). This demonstrates the capacity of the K-GG antibody beads to enrich high amounts of modified peptide, and is also, to our knowledge, the highest number of K-GG peptides ever identified in a single shot LC-MS/MS analysis. DIA analysis of both untreated and MG132 treated samples returned much higher numbers of modified peptides than DDA analysis on the same instrument, demonstrating the value of nDIA in analysis of post translationally modified peptides. This study also yielded over 33,000 novel sites of ubiquitin modification compared to known sites in the PhosphoSitePlus database, massively expanding the number of known ubiquitination sites and providing a rich resource for future studies of protein ubiquitination (**Figure 2G**).

Previous work has demonstrated the utility of the Orbitrap Astral MS instrument in phosphoproteomics experiments [41]. This study confirmed those earlier findings that DIA on Orbitrap Astral MS allows significant gains in depth of phosphopeptide coverage using Fe-IMAC enrichment (**Figure 3A**, **3B**). The vast majority of sites identified from IMAC enriched samples were phosphoserine and phosphothreonine (∼98%) as described previously [19]. To achieve more comprehensive coverage of the tyrosine phosphoproteome a phosphotyrosine (pY) specific enrichment is needed, such as the PTMScan phosphotyrosine pY-1000 kit. Indeed, this enrichment allowed identification of hundreds to thousands of phosphotyrosine sites across samples (**Figure 2C**, **2D**), and the use of Orbitrap Astral MS provided a large increase in pY peptides identified compared to the older generation Orbitrap Fusion Lumos MS instrument, with up to 33-fold more pY peptides identified with Orbitrap Astral nDIA than Orbitrap Fusion Lumos DDA. As with MG132 for ubiquitination analysis, the tyrosine phosphatase inhibitor pervanadate was used to treat Jurkat cells and create a maximally phosphorylated sample (**Figure 3E**). Pervanadate treatment allowed identification and quantification of over 10,000 phosphotyrosine sites with Orbitrap Astral DIA, compared to fewer than 1,000 sites with untreated Jurkat cells (**Figure 3F**). This again demonstrates the capacity for peptide enrichment of the beads, as well as the gains in depth of coverage that can be made by employing nDIA on the Orbitrap Astral MS. 10,000 phosphotyrosine sites represent one of the larger single shot datasets collected to date, and overall, the phosphotyrosine data in this study contributed more than 10% new sites to the PhosphoSitePlus database (**Figure 3G**), even though pY data has been curated into that database for over 20 years.

Acetylation profiling was similarly improved with nDIA on Orbitrap Astral MS, with nearly 3-fold more AcK peptides identified and quantified with Orbitrap Astral DIA-HBS compared to Orbitrap Fusion Lumos DDA, and nearly 26% additional sites not previously in the PhosphoSitePlus database (Figure 4A, 4B). The gains for succinyl lysine and mono-methyl arginine were more subtle, however in both cases nDIA analysis on Orbitrap Astral MS outperformed DDA on Orbitrap Fusion Lumos MS despite a 2X longer gradient used for the DDA Orbitrap Fusion Lumos MS analysis (**Figure 4C**, **4E**). Again, as with other modifications, the nDIA profiling on Orbitrap Astral MS contributed new sites to the PhosphoSitePlus database, with more than 2X new sites for succinylation (110% novel, Figure 4D), and 18% new sites for mono-methyl arginine (**Figure 4F**).

For each PTM profiled in the study, an unenriched sample was run in parallel using nDIA on Orbitrap Astral MS. This served two purposes: To evaluate the ability of the Orbitrap Astral MS, given its speed and sensitivity, to identify PTM peptides without the need for enrichment, and to verify that DIA searching is not providing spurious identification of PTM peptides (Figure 5A-5G). In all cases, there were very few modified peptide identifications in the unenriched samples, while enrichment allowed identification and quantification of thousands to tens of thousands of modified peptides, highlighting the need to enrich samples to effectively profile PTMs. Likewise, the low number of PTM peptide identifications in the unenriched samples demonstrates that the DIA software is not providing artificial identification of PTM peptides in a sample where they have not been specifically enriched.

Enrichment methods such as the PTMScan HS technology circumvent the challenge of identifying low abundance post translationally modified peptides through immunoaffinity enrichment of peptides harboring specific class of PTMs including but not limited to lysine ubiquitination, tyrosine phosphorylation, lysine acetylation, lysine succinylation, and arginine mono-methylation. The Astral Orbitrap MS system represents a major leap forward in the speed, sensitivity, and depth of coverage achievable for bottom-up proteomics, owing to its nearly lossless ion transfer ability, combined with ∼200-Hz MS/MS acquisition rates at high resolving power [28, 30, 31]. Combining PTMScan sample preparation with the Astral Orbitrap MS instrument operated under narrow window data-independent acquisition (nDIA) allows unprecedented access to the PTM landscape in cells and tissues. For all PTMs tested in this study, identification of PTM-containing peptides with Orbitrap Astral nDIA far outpaced the previous generation Orbitrap Fusion Lumos MS instrument. These data also represent a major improvement in the catalog of known PTMs in human cells and mouse tissue, with thousands of unique PTM sites identified that are not currently in the PhosphoSitePlus database. The protocol detailed in this study will provide a framework for future analysis of protein post-translational modifications in the study of cellular signaling, disease biology, and target and biomarker discovery.

## Abbreviations

PTM: post-translational modifications
DDA: Data Dependent Acquisition
nDIA: Narrow Window Data Independent Acquisition
IMAC: Immobilized Metal Affinity Chromatography
PTMScan HS: post translational modification scan high specificity
HCT116: human colorectal carcinoma cell
Jurkat: human T-cell leukemia cell
DMSO: dimethyl sulfoxide
DDT: Dithiothreitol; SDS-PAGE, sodium dodecyl sulfate polyacrylamide gel electrophoresis
HRP: horseradish peroxidase
HEPES: 4-(2-hydroxyethyl)-1-piperazineethanesulfonic acid
IAP: Immunoaffinity purification
TFA: trifluoroacetic acid
MeCN: acetonitrile
FA: formic acid
LFS: Library Free Search
HBS: Hybrid Search.

## Data Availability

The mass spectrometry proteomics data have been deposited to the ProteomeXchange Consortium via the PRIDE [42] partner repository with the dataset identifier PXD065579

## Acknowledgments

We thank Michael J. Comb and Cell Signaling Technology for funding this research project.

## Author Contribution

M.K., A.P.P., J.C.S., M.P.S., S.A.B. **Conceptualization and study design**: M.K., A.P.P., B.M.Z., S.S., J.M.R., A.J.N., B.L., T.P.H.; **Methodology**: M.K., A.P.P., S.S., J.M.R., B.Z., S.L., B.L., T.P.H.; **Data acquisition and analysis**: M.K., S.S., B.Z., S.L., M.P.S., **Visualization**: M.K., A.P.P., J.C.S., M.P.S.; **Writing–original draft**: M.K., A.P.P., B.M.Z., S.S., J.M.R., A.J.N., B.Z., S.L., J.C.S., B.L., T.P.H., M.P.S., S.A.B. **Writing reviewing and editing**

## Conflict of interest

M.K., A.P.P., B.M.Z., S.S., J.M.R., A.J.N., B.Z., S.L., J.C.S., M.P.S., and S.A.B. are employees of Cell Signaling Technology. B.L., and T.P.H., are employees of Thermo Fisher Scientific.

